# Microbiome engineering could select for more virulent pathogens

**DOI:** 10.1101/027854

**Authors:** Luke McNally, Pedro F. Vale, Sam P. Brown

## Abstract

Recent insights into the human microbiome offer the hope of manipulating its composition to help fight infectious diseases^1–7^. While this strategy has shown huge potential, its consequences for pathogen evolution have not been explored. Here we show that manipulating the microbiome to increase the competition that pathogens face could lead to the evolution of increased production of virulence factors that pathogens use to combat commensals, an evolutionary response that can increase total disease induced mortality in the long-term. However, if treatment with microbiome engineering is sufficiently aggressive this evolutionary response can be avoided and the pathogen eradicated. Furthermore, we show that using damage limitation therapies^8^ (e.g. antivirulence and anti-inflammatory drugs) in combination with microbiome manipulation increases the potential for pathogen eradication. While manipulating our microbiota offers a promising alternative to antibiotics, our results show that these treatments must be designed with careful consideration of the potential evolutionary responses of target pathogens.

## Introduction

The antibiotic resistance crisis has led to the search for alternative ways to treat bacterial infections. One of the most promising forms of novel treatment strategies is to modify the commensal microbiome in order to either prevent pathogen colonisation or remove the pathogen once it has colonised^9^. The central idea behind this approach is to introduce bacteria that can competitively suppress the pathogen. This strategy has already demonstrated its promise with ‘faecal transplant’ therapies showing high success rates in curing recurrent *Clostridium difficile* infections that are recalcitrant to antibiotic treatment^1,2^. The remarkable success of microbiome manipulation in treating *C. difficile* infections suggests that this approach may be viable as a treatment for other infections, with suggestions that this approach could be used to prevent *Staphylococcus aureus* nasal carriage^3,4^, to prevent dental caries^5^, and to prevent infections in plants to improve crop yields^6,7^.

While microbiome manipulation may at first appear to be a robust way to treat or prevent infection, the amazing ability of bacterial pathogens to adapt to both antibiotic treatments^10^ and vaccines^11^ shows that we must consider the evolutionary consequences of any novel treatment strategies. So, how could pathogens evolve resistance to microbiome manipulation? Overcoming competition from commensals is not a new challenge for pathogens and they have evolved an array of weapons to help them remove commensal competitors and colonise their host, either by directly killing competitors or provoking host inflammation to which they are resistant^12-14^.

Importantly, these weapons are often direct causes of virulence in the host. Examples in human pathogens include suicidal invasion of gut tissue in order to provoke inflammation in *Salmonella enterica* serovar Typhimurium^15,16^; release of shiga toxin encoding phage by shigatoxinagenic *Escherichia coli*, which removes competitors both directly and via inflammation^13,17^; release of the toxin pyocyanin by *Pseudomonas aeruginosa*^18,19^; provocation of host immune responses by *Haemophilus influenzae*^20^; and release of the toxin TcdA by *C. difficile* causing inflammation that may clear commensals^16,21-23^. Increasing the strength of competition that these pathogens face from commensals could create selection for increased expression of these weapons, potentially leading to the evolution of increased virulence. Here we use a multi-level model of both within-host competition between a pathogen and commensals and the epidemiological spread of the pathogen to examine the evolutionary and epidemiological consequences of the use of microbiome manipulation as a treatment strategy.

## Results and Discussion

We consider a scenario where a pathogen competes with an introduced therapeutic commensal bacterium at the disease site (Fig. 1a). We assume for simplicity that both species have the same basal per capita growth rate *r* (though relaxing this assumption does not qualitatively affect our conclusions). The commensal competitor (at frequency 1 - *P*) reduces the growth rate of the pathogen by amount *a*(1 - *P*). The pathogen (at frequency *P*) produces amount *v* of a virulence factor, which reduces the commensal’s growth rate (either directly or via interactions with the immune system) by amount *bvP*, where *b* is the sensitivity of the commensal to the effects of the virulence factor. The pathogen pays a growth rate cost *c* per unit virulence factor expression. Analysing this system (see methods) we see that the pathogen free equilibrium (*P* = 0) is always locally stable, while pathogen dominance (*P* = 1) is locally stable whenever *b* > *c* (i.e. as long as commensal sensitivity to the virulence factor is greater than the cost of it’s expression, Fig. 1b). There is also an internal unstable equilibrium (Fig. 1b) given by

**Fig. 1.**
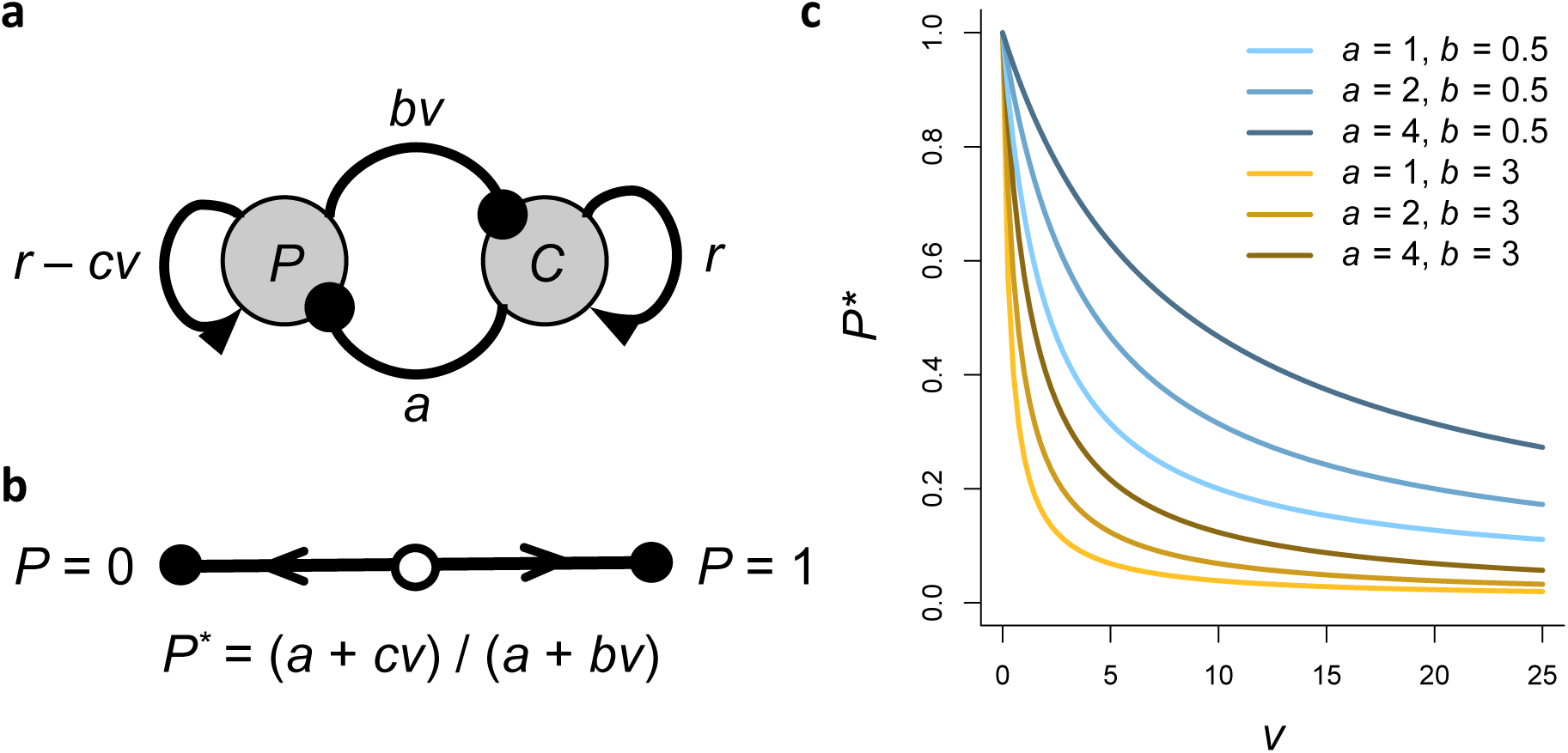
Within-host competition between pathogen and commensals. (a) Schematic of the within host model described in equation 1. Triangular arrowheads indicate growth of each population, while circular arrowheads indicate suppressive effects, (b) Illustration of within host dynamics. Stable equilibria are illustrated by solid points, with the threshold pathogen frequency (unstable equilibrium) is illustrated by the open circle, (c) Behaviour of pathogen frequency threshold. The frequency threshold is plotted as a function of the suppressive effect of the commensal, *a*, the sensitivity of the commensal to the virulence factor, *b*, and pathogen investment in virulence factor production, *v.* Parameters values are *c* = 0.02.

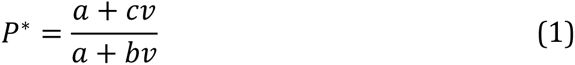

If the frequency of the pathogen goes above this threshold then the pathogen domination equilibrium is reached (*P* = 1), while if the pathogen frequency goes below this threshold then the pathogen free equilibrium is reached (*P* = 0) (Fig. 1b). This threshold will therefore decide how likely it is that a preventative treatment stops the pathogen invading (stops transition from *P* = 0 to *P* = 1), and how likely it is for responsive treatment to clear an infection (cause transition from *P* = 1 to *P* = 0). This threshold pathogen frequency becomes higher (making it easier to clear the pathogen) with increasing suppression by the commensal (*a*) and increasing costs of virulence factor expression (*c*), while the threshold is lowered (making it more difficult to clear the pathogen) by increasing sensitivity of the commensal (*b*) and increasing virulence factor expression (*v*)(Fig. 1c).

To model the evolutionary and epidemiological consequences of this competition between pathogens and introduced commensals we use our within-host model to derive an epidemiological model for the spread of strains with different levels of investment in virulence factor production. Here we present a model for responsive treatment aimed at clearing a pathogen once infection has been established (such as the case of faecal transplant therapy in reponse to *C. difficile* infection), but we find qualitatively similar results for a model of preventative treatment by modifying the microbiome of healthy individuals (see supplementary information). The ‘reproductive number’ (number of secondary infections caused when the pathogen is rare) of a pathogen strain of virulence *v* is

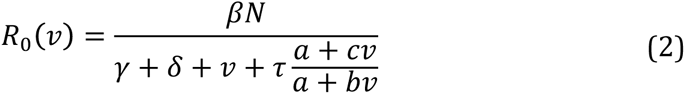

where *N*, is the host population density, *β* is a transmission constant, *γ* is the baseline clearance rate, *δ* is the baseline disease induced mortality, and *τ* is the treatment rate. We can calculate the evolutionarily stable (ES) level of virulence factor production (*v*^*^) by maximising *R*_0_(*v*), giving

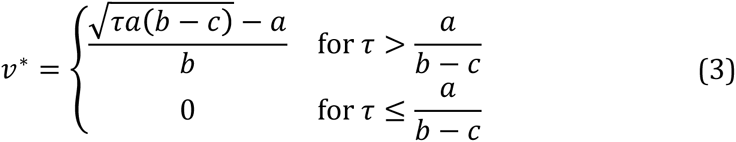

We can see from equation 3 that increasing rates of treatment with microbiome manipulation (higher *τ*) select for increases in virulence factor production (Fig. 2a). However, while microbiome manipulation increases selection for virulence factor production, it simultaneously reduces the prevalence of the pathogen at equilibrium by increasing the infection clearance rate, and can even lead to pathogen eradication at high treatment rates (Fig. 2b). These conflicting effects of increasing pathogen virulence and decreasing pathogen prevalence leads to a humped relationship between treatment rate and the total disease induced mortality at equilibrium (Fig. 2c), meaning that despite eradicating the pathogen at high treatment rates, microbiome manipulation can actually lead to increased disease induced mortality if treatment is not sufficiently aggressive.

**Fig. 2.**
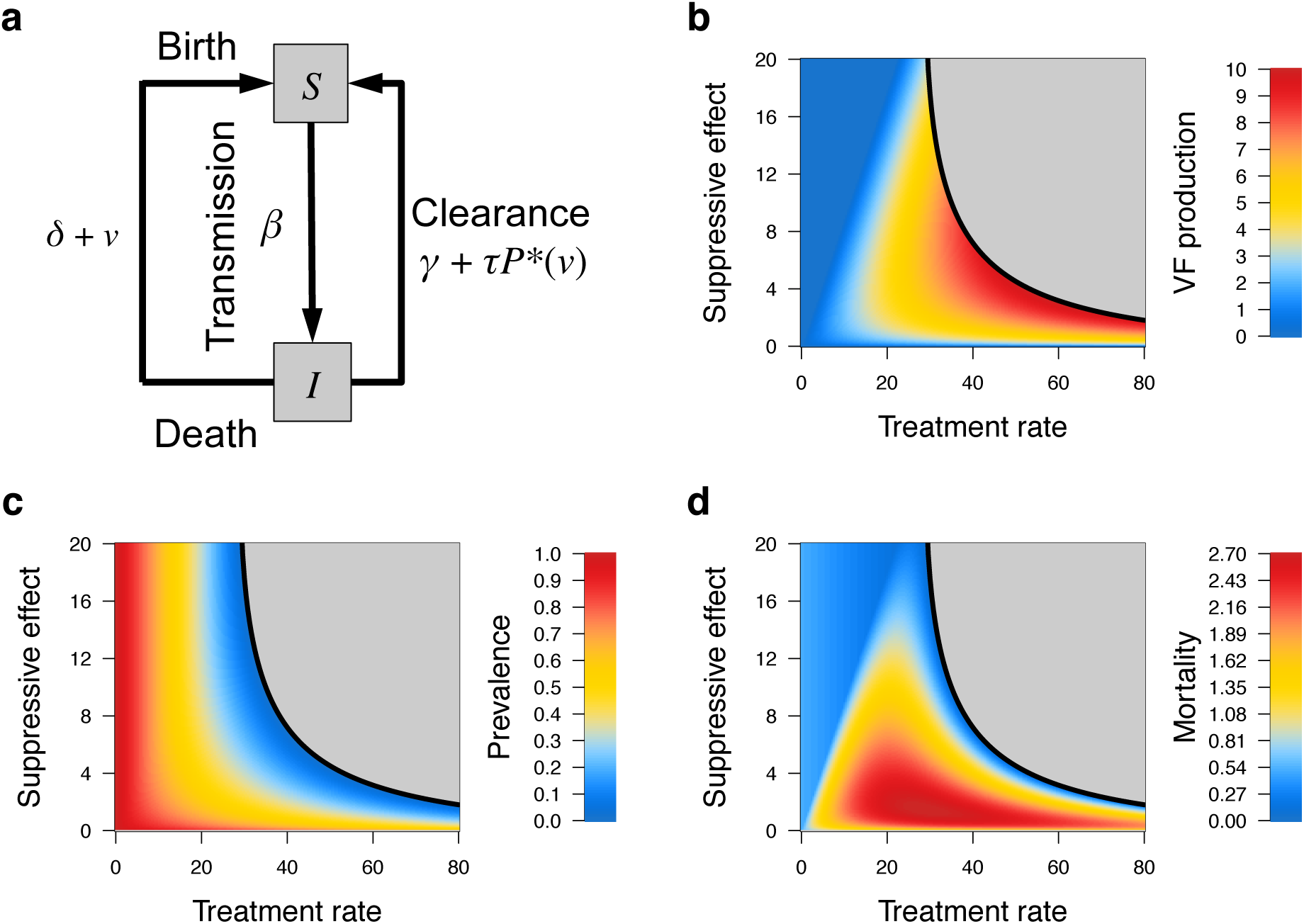
Evolutionary and epidemiological consequences of manipulating the microbiome. (a) Schematic of the epidemiological model described in equation 2. (b-d) Plotted are the predicted VF production, *v*^*^ (b), pathogen prevalence, *Q*^*^ (c), and the total disease induced mortality at evolutionary equilibrium, (*v*^*^ + *δ*)*Q*^*^ (d), as a function of the suppressive effect of the commensal, *a*, and the treatment rate, *τ.* The grey area indicates where the pathogen is eradicated despite its evolution. Parameter values for (b-d) are *b* = 1, *c* = 0.05, *β* = 3, *N* = 10, *γ* = 0.5, *δ* = 0.5.

Our analytical model assumes that the pathogen will evolve to its optimal level of virulence factor production in the face of microbiome manipulation. However, if the pathogen cannot evolve higher levels of virulence factor production quickly enough it could be eradicated by treatment before it reaches its optimal virulence. To explore this scenario we built a stochastic simulation model of the evolution of virulence factor production under treatment with microbiome manipulation. Our simulations show that the evolution of increased virulence factor production acts as a form of ‘evolutionary rescue’^24^ for the pathogen – if the pathogen has a sufficiently high mutational supply it can avoid eradication by evolving increased virulence (Fig. 3). This evolutionary rescue becomes less likely with higher treatment rates as the pathogen population size declines rapidly, reducing its mutational supply (Fig. 3).

**Fig. 3.**
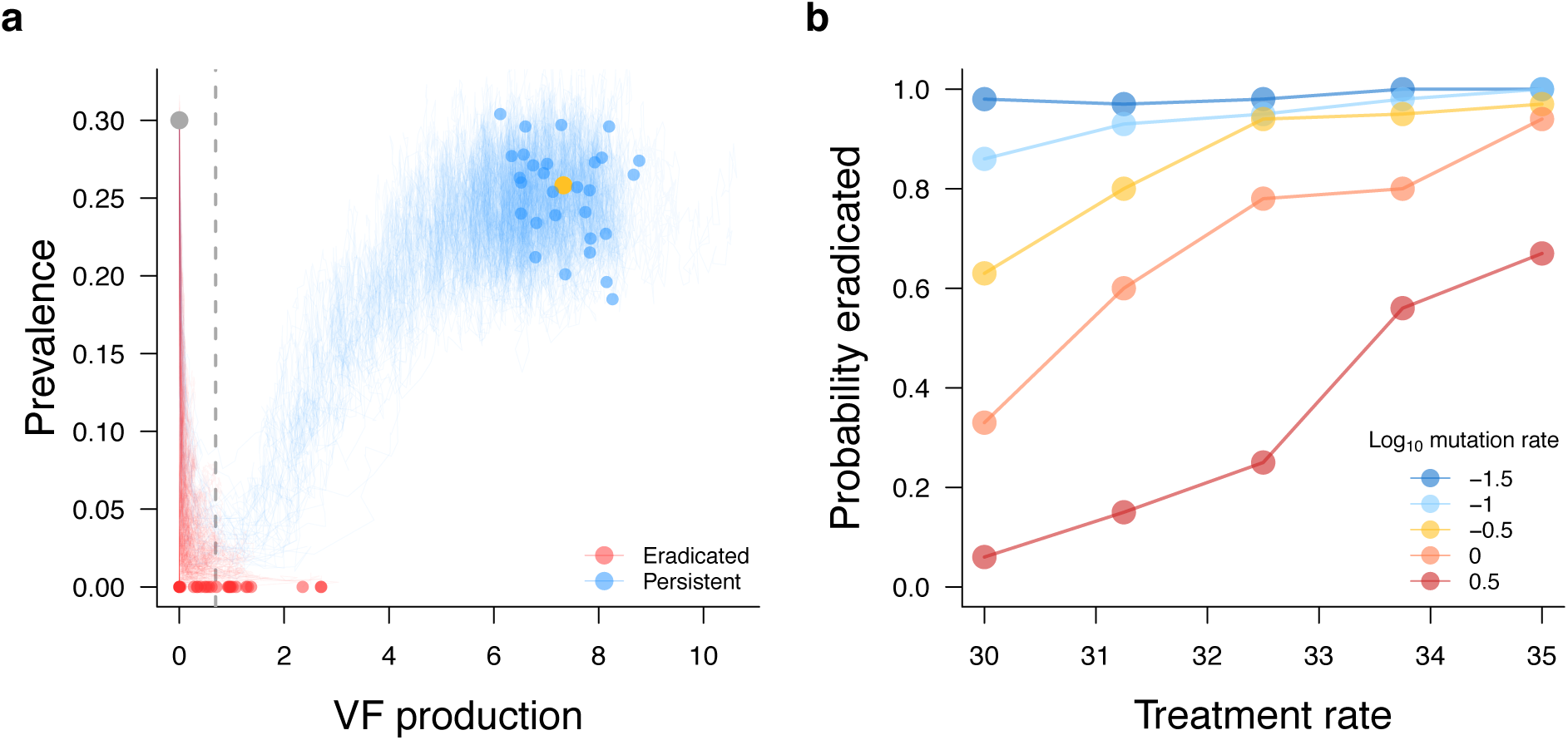
Evolutionary rescue of pathogens via evolution of increased virulence. (a) Plotted are 100 sample trajectories of our stochastic simulation of our model. The grey dot indicates the starting point of the simulation and the yellow dot indicates the equilibrium predicted by our analytical model. Blue and red lines and points indicate the evolutionary trajectories and finishing points of replicate simulations, blue where the pathogen persisted and red where the pathogen was eradicated. The grey dashed line indicates the minimum level of VF production where the pathogen persists in a deterministic system. See Movie S1 for a video of the dynamics. (b) Plotted is the probability that the pathogen is eradicated as a function of the treatment rate and rate of supply of mutations affecting VF production. Each point comes from 100 independent stochastic simulations. Parameter values are *N* = 1000, *β* = 0.03, *a* = 5, *b* = 1, *γ* = 0.5, *δ* = 0.5, *σ_μ_* = 1 for (a-b), and *μ* = 1, *τ* = 32 for (a).

If sufficiently high treatment intensity with microbiome manipulation cannot be achieved pathogens can undergo evolutionary rescue via increased virulence factor production, which can lead to increased disease induced mortality. However, microbiome manipulation could be used with other co-therapies to increase its efficacy. One approach that may allow sufficient increase in efficacy to avoid this outcome is to combine microbiome manipulation with damage limitation therapies^8^. Damage limitation therapies limit the pathogenesis of infection without directly killing pathogens, by either inhibiting the production or action of pathogen virulence factors (anti-virulence drugs^8,25^), or by increasing the host’s capacity to limit and repair tissue damage from both the infection and its own immune response (pro-tolerance drugs^8,26^). These drugs thus will either directly inhibit the effects of the pathogen’s virulence factors (anti-virulence) or stop the induced inflammation (pro-tolerance), and could therefore provide a powerful synergy with microbiome manipulation by reducing the pathogen’s ability to attack introduced commensals. We modified our model to consider a scenario where a damage limitation therapy (of treatment intensity *x*) is given alongside manipulation of the microbiome. This modification of our model shows that using damage limitation therapy in combination with microbiome manipulation expands the range of conditions where the pathogen can be eradicated, and can offset increases in mortality owing to increased selection for virulence factor production by decreasing the virulence factors effects on the host (Fig. 4).

**Fig. 4.**
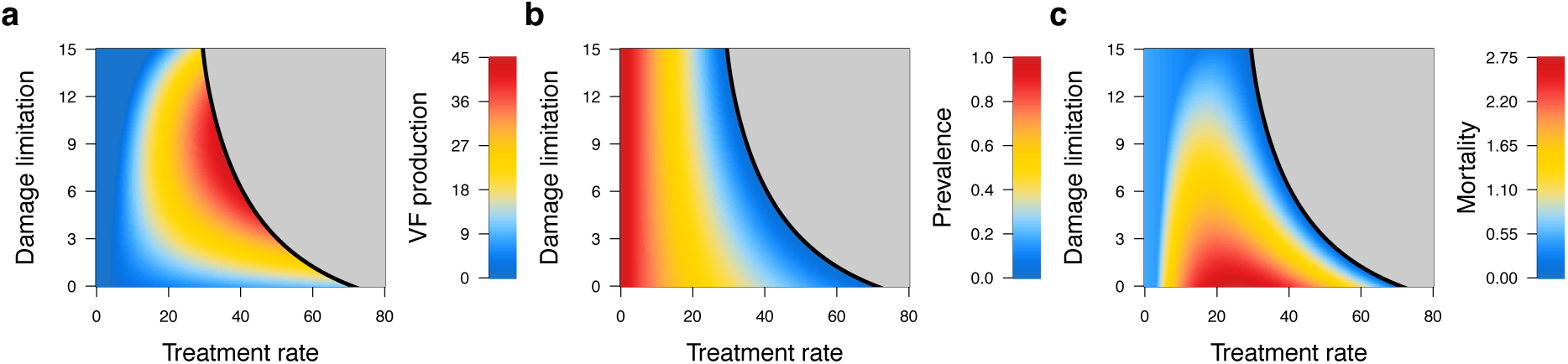
Combining microbiome manipulation and damage limitation. (a-c) Plotted are the predicted VF production, *v*^*^ (a), pathogen prevalence, *Q*^*^ (b), and the total disease induced mortality at evolutionary equilibrium, (*v*^*^/(1 + *x*) + *δ*)*Q*^*^, as a function of the efficacy of damage limitation, *x*, and treatment rate, *τ*. The grey area indicates where the pathogen is eradicated despite its evolution. Parameter values for (a-c) are *a* = 2, *b* = 1, *c* = 0.05, *β* = 3, *N* = 10, *γ* = 0.5, *δ* = 0.5.

Advances in microbiome sequencing and techniques to discover interactions among microbes from sequence data offer the hope of allowing us to engineer the human microbiota to repel or clear invading pathogens^27^. While this is a highly promising therapeutic avenue, just as with other forms of antimicrobial treatment there is the potential for pathogens to evolve resistance to our interventions. Our results show that one potential pathogen response to microbiome manipulation is to upregulate its arsenal of virulence factors that it uses to clear commensals. This leads to a scenario analogous to the ‘double-edged sword’ effects of antimicrobial drug dosing^28^, where higher doses of drugs maximise selection for resistance, but also reduce the supply of resistance mutations, reduce prevalence, and increase the chances of pathogen eradication. The overarching lesson from this problem has been that we need to carefully consider the evolutionary responses of pathogens when designing treatment strategies^28^. Given that an expected evolutionary response of pathogens to increases in competition is to increase their virulence, it is critical to heed this lesson when designing manipulations of the microbiome.

Where microbiome manipulation is not efficacious enough to achieve pathogen eradication our results suggest the use of co-therapies that reduce the impact of pathogen virulence factors. These ‘damage limitation’ therapies are the subject of much current interest as they are expected to show less resistance evolution than traditional antimicrobials^8,25^. Our results suggest that by reducing the impact of pathogen virulence factors on commensals, damage limitation therapies could show a strong synergy with microbiome manipulation and help achieve pathogen eradication.

Microbiome manipulation has already been successfully deployed to treat recurrent *C. difficile* infections via faecal transplant^1,2^, and the ecological mechanisms underlying this therapy are increasingly well understood, offering hope of increased efficacy through more precise treatment in the future^27^. However, current faecal transplant therapy has a failure rate of approximately 8%-19%^1,2^. We would suggest that monitoring of toxin production of *C. difficile* cases before and after faecal transplant treatments, as well as comparison between strains from successful and unsuccessful treatments, could help detect selection for increased toxin production as a result of treatment. In addition, there are several anti-toxin vaccines for *C. difficile* in various stages of clinical trial^29^, as well as a newly discovered anti-virulence drug^30^. Given that the *C. difficile* toxins TcdA and TcdB induce inflammation that clears competitors^16,21-23^, our results suggest that combination of these anti-toxin vaccines with microbiome manipulations could help effectively eradicate *C. difficile* infection and prevent evolutionary increases in it’s virulence.

Bacterial species have been competing throughout a long evolutionary history and have evolved highly efficient weapons to target their competitors. Increasing the intensity of bacterial warfare within us requires consideration of its evolutionary consequences, as human health is often a bystander casualty in this conflict.

## Methods

### Within-host model

We model the dynamics of the pathogen and introduced commensal using the replicator equation^31^, with the dynamics of the within-host frequency of the pathogen, *P*, given by

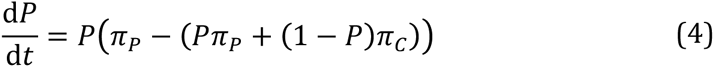

where *π_P_* = *r* − *cv* − *a*( 1 − *P*) and *π_c_* = *r* − *vbP* are the growth rates of the pathogen and commensal, respectively. Evaluating *P* at *dP/dt* = 0, and assuming *b* > *c*, we find stable equilibria at *P* = 0 and *P* = 1, and an unstable equilibrium given in equation 1.

### Epidemiological model

We use the simplest scenario of a susceptible-infected epidemiological model to model the spread of strains with different levels of investment in virulence factor production. We assume that host deaths are immediately replaced by births and that treatment-induced clearance is proportional to the location of the threshold for invasion by the introduced commensal^12,32^. Using these assumptions the epidemiological dynamics are given by

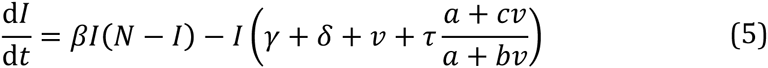

where *N*, is the host population density, *I* is the density of infected hosts, *S* = *N* − *I* is the density of susceptible hosts, *β* is a transmission constant, *γ* is the infection clearance rate, *δ* is the baseline host death rate, *τ* is the treatment rate, and virulence factor production increases the host death rate by *v*. From this we can write the ‘reproductive number’ (number of secondary infections caused when rare)^33^ by evaluating the ratio of transmission and loss rates when the pathogen is rare (*I* → 0) giving equation 2. The ES level of virulence factor production (*v*^*^) by finding the value of *v* that maximises *R*_0_(*v*), giving equation 3, which shows that the ES virulence factor production (*v*^*^) increases with increased treatment rates (*τ*).

To evaluate the impact of microbiome manipulation on the long-term epidemiology of the pathogen we evaluate the pathogen *R*_0_ at evolutionary equilibrium (*v* = *v*^*^), giving

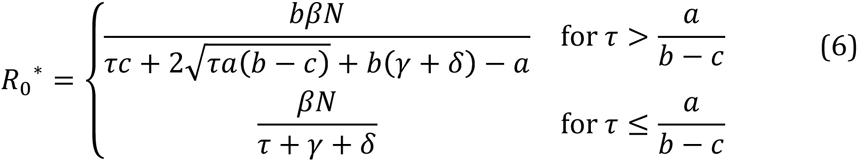

which must be greater than 1 for the pathogen to avoid eradication. This R0 corresponds to an equilibrium pathogen prevalence of

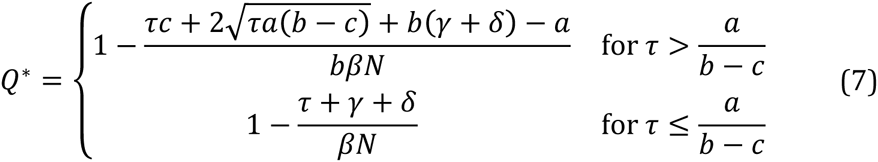

We can see from equation 7 that pathogen prevalence, *Q*^*^, declines with increasing rates of treatment with microbiome manipulation, *τ*, with the pathogen being eradicated at sufficiently high treatment rates.

Given that introducing competitors in the microbiome selects for higher virulence but also reduces the prevalence of infection what is the impact of microbiome manipulation on the total disease induced mortality? The total disease induced mortality at evolutionary equilibrium is given by *M*^*^ = (*v*^*^ + *δ*) *Q*^*^, which is

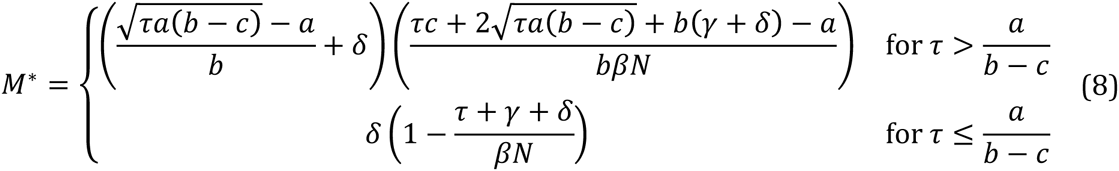

Analysing equation 8, and from Fig. 2d, we can see that the then the total disease induced mortality can increase with *τ*, though must decline with *τ* at higher levels as the pathogen approaches eradication. This result shows that the evolutionary risks of microbiome manipulation must be carefully considered – if treatment is not sufficiently aggressive it may result in increased disease induced mortality owing to evolutionary increases in virulence.

### Combining microbiome manipulation and damage limitation

We consider a damage limitation therapy, of treatment intensity *x*, used in combination with microbiome manipulation. By disrupting the effects of the virulence factor the damage limitation therapy reduces both the pathogen’s effect on introduced commensals and damage to the host. Weighting these effects by damage limitation efficacy the pathogen *R*_0_ is now

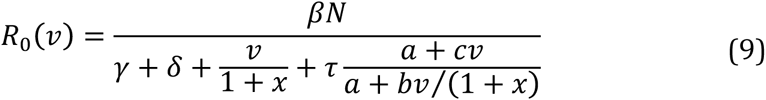

and the ES virulence is

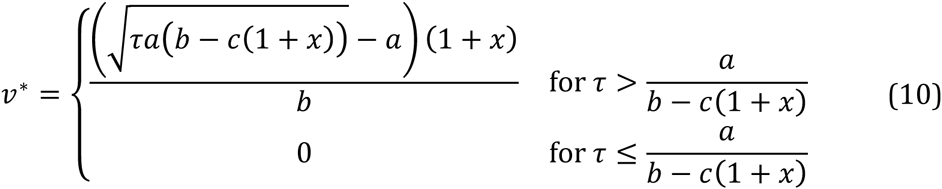

Here we can see that damage limitation therapy has a non-monotonic (humped) effect on ES virulence. Again we evaluate the pathogen Ro at evolutionary equilibrium (*v* = *v*^*^), giving

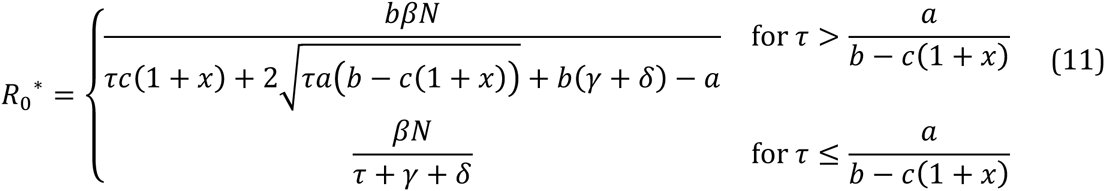

and corresponding to an equilibrium prevalence of

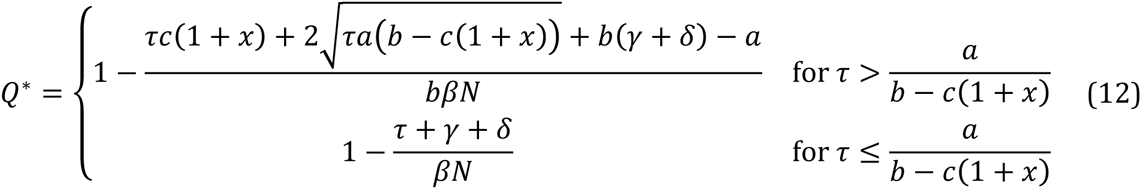

which decreases with *x* whenever *τ* > *a/*(*b* − *c*( 1 + *x*)), meaning damage limitation therapy helps reduced pathogen prevalence by reducing the pathogen’s ability to attack the commensal competitors via its virulence factors. The total disease induced mortality at evolutionary equilibrium is given by

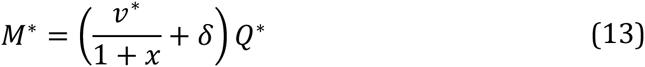

the full expression for which can be calculated using equations 10 and 12. Note that though damage limitation increases the ES virulence factor expression, this is offset by damage limitation reducing the efficacy of the virulence factor; meaning that the total disease induced mortality decreases with damage limitation therapy. Note however, that this only holds as long as damage limitation therapy is continued to be used, ceasing use while the pathogen is still at appreciable prevalence may increase mortality rates^8^.

### Stochastic simulations of evolutionary rescue

To simulate the stochastic dynamics of the evolution of virulence factor expression in response to microbiome manipulation treatment we simulated our epidemiological model using the Gillespie algorithm^34^. Our model consists of a finite population of *N* hosts, which can be infected with strains with differing levels of virulence factor production. The rates of transmission, clearance of infection, and host death (immediately replaced by birth of an uninfected host) are calculated for each infection as per equation 5. In addition to these processes the strain in each infection can mutate with mutation rate *μ*, in which case its virulence factor expression has a value added to it from a normal distribution with mean 0 and variance *σ_μ_*.

## Acknowledgements

We thank Roman Popat, Laura Pollitt, and Tom Little for comments on previous versions of this manuscript. LM and SPB are supported by the Human Frontier Science Programme (RGP0011/2014). PV is supported by a Chancellor’s fellowship from the University of Edinburgh and a Society in Science – Branco Weiss fellowship (http://www.society-in-science.org - ETH Zürich). All authors were also supported by the Wellcome Trust supported CIIE (grant ref. WT095831).

